# Cellular electron tomography of the apical complex in the apicomplexan parasite *Eimeria tenella* shows a highly organised gateway for regulated secretion

**DOI:** 10.1101/2021.06.17.448283

**Authors:** Alana Burrell, Virginia Marugan-Hernandez, Flavia Moreira-Leite, David J P Ferguson, Fiona M Tomley, Sue Vaughan

**Author notes:** Joint corresponding author.

## Abstract

The apical complex of apicomplexan parasites is essential for host cell invasion and intracellular survival and as the site of regulated exocytosis from specialised secretory organelles called rhoptries and micronemes. Despite its importance, there is little data on the three-dimensional organisation and quantification of these organelles within the apical complex or how they are trafficked to this specialised region of plasma membrane for exocytosis. In coccidian apicomplexans there is an additional tubulin-containing hollow barrel structure, the conoid, which provides a structural gateway for this specialised secretion. Using a combination of cellular electron tomography and serial block face-scanning electron microscopy (SBF-SEM) we have reconstructed the entire apical end of *Eimeria tenella* sporozoites. We discovered that conoid fibre number varied, but there was a fixed spacing between fibres, leading to conoids of different sizes. Associated apical structures varied in size to accommodate a larger or smaller conoid diameter. However, the number of subpellicular microtubules on the apical polar ring surrounding the conoid did not vary, suggesting a control of apical complex size. We quantified the number and location of rhoptries and micronemes within cells and show a highly organised gateway for trafficking and docking of rhoptries, micronemes and vesicles within the conoid around a set of intra-conoidal microtubules. Finally, we provide ultrastructural evidence for fusion of rhoptries directly through the parasite plasma membrane early in infection and the presence of a pore in the parasitophorous vacuole membrane, providing a structural explanation for how rhoptry proteins (ROPs) may be trafficked between the parasite and the host cytoplasm.

**Significance:** Apicomplexan parasites cause a wide range of human and animal diseases. The apical complex is essential for motility, host cell invasion and intracellular survival within a specialised vacuole called the parasitophorous vacuole. We know that molecules important for all of these processes are secreted from the apical complex via a set of secretory organelles and there is even evidence that some parasite molecules can enter the host cell from the parasitohorous vacuole, but there is little understanding of exactly how this occurs. Here we have used three dimensional electron microscopy to reconstruct the entire apical end of the parasite and whole individual parasites. Our results provide important insights into the structural organisation and mechanisms for delivery of parasite molecules via this important area of the cell.

## Introduction

A defining feature of apicomplexan parasites is the apical complex after which the phylum is named. This comprises an apical polar ring to which are attached a set of helically arrayed subpellicular microtubules, and two types of specialised secretory apical organelles called micronemes and rhoptries that are part of the parasite endomembrane system. The Coccidia sub-class of Apicomplexa, which includes the genera *Eimeria, Neospora, Sarcocystis* and *Toxoplasma*, also possess a conoid, an apical cone-like hollow structure composed of tubulin-containing fibres (Morrissette and Sibley, 2002) associated with two pre-conoidal rings and a pair of intra-conoidal microtubules (Nichols and Chiappino, 1987). Transmission electron microscopy analysis in *Toxoplasma gondii* estimated the conoid to contain ~14 tubulin fibres (Hu et al., 2002), but there was a level of uncertainty in this number due to inherent difficulties in analysing such a complex structure using two-dimensional methods. The two pre-conoidal rings are located at the apical tip of the cell, above the conoid and just under the plasma membrane, and these rings move together with the mobile conoid when it is extended in *T. gondii*, although their precise role is not understood (Nichols and Chiappino, 1987; Katris et al., 2014).

Co-ordinated exocytosis of apical secretory organelles during host cell invasion is essential not only for parasite entry into the host cell, but also for intracellular survival. Micronemes secrete adhesin complexes (MICs) onto the parasite surface where they span the parasite plasma membrane (PM) and connect to a parasite actinomyosin-based motility system, the glideosome, which occupies a space in the parasite pellicle between the PM and the inner membrane complex for parasite motility in extracellular parasites (Frénal et al., 2017). Surface exposure of MICs is essential for attachment to host cells, gliding motility, migration across tissues and for the formation of a moving junction (MJ) between the parasite and the host cell, during invasion (Carruthers and Tomley, 2008). A complex of rhoptry neck proteins (RONs) are inserted into the host cell PM and act as cognate receptors for the MIC protein AMA1, which is itself anchored to the parasite apical surface after secretion. The AMA1-RON2 interaction is critical for the formation of a stable moving junction (Lamarque et al., 2011) and the RON complex recruits host cytoskeletal structures to the cytosolic face of the host PM, to solidify host-parasite bridging as invasion progresses (Guérin et al., 2017). The MJ also sieves and removes GPI-anchored proteins from the host PM so that the newly formed parasitophorous vacuole (PV) can resist endosome fusion and lysosomal destruction (Mordue et al., 1999). Secreted rhoptry bulb proteins (ROPs) and membranous material contribute to the formation and remodelling of the PV and its membrane and some ROPs traffic beyond the PV into the host cell cytoplasm (Bradley and Sibley, 2007).

Secretion of both rhoptry and microneme proteins occurs from the apical complex, but the precise organisation of these trafficking events is not well understood. MIC secretion from micronemes occurs from the apical pole when parasites contact host cells (Frénal et al., 2017; Bumstead and Tomley, 2000). This contact activates a parasite cGMP signalling pathway with two effector arms. Inositol triphosphate (IP3) stimulates intracellular calcium release activating calcium-dependent protein kinases (CDPKs) at the parasite apex. CDPKs then phosphorylate proteins involved in microneme exocytosis and in the activation of the glideosome (Dunn et al., 1996). Diacylglycerol (DAG) is converted to phosphatic acid at the inner leaflet of the parasite PM and sensed by a pleckstrin homology domain protein (APH) that lies on the surface of apical micronemes (Bullen et al., 2019). Despite these advances in understanding the signalling pathway, the actual mechanism of microneme exocytosis remains enigmatic. RON/ROP secretion from rhoptries also occurs at the apical pole, but apart from needing previous MIC discharge (Kessler et al., 2008) little is known about the signalling pathway triggering rhoptry discharge. Apical positioning of the rhoptry seems to be prerequisite for RON/ROP exocytosis (Frénal et al., 2013) and it is not possible to trigger RON/ROP secretion in extracellular parasites. However invasion is not essential; if actin polymerisation (and invasion) are disrupted by treatment with cytochalasin D parasites are still able to attach, form a tight junction with the host PM and discharge rhoptry content into nascent parasitophorous ‘evacuoles’, which can be readily detected with anti-ROP antibodies (Håkansson et al., 2001). Indeed it appears that in normal *in vitro* and *in vivo* infections a proportion of host cells are injected with rhoptry proteins without becoming infected; the reason for this is unknown (Koshy et al., 2012). After discharge, rhoptries are occasionally seen as empty sacs (Lebrun et al., 2005) and in intracellular parasites one or two rhoptries may be seen with their necks extending into the conoid reaching the apical PM, suggesting they are anchored here and ready for secretion (Paredes-Santos et al., 2012). The two intra-conoidal microtubules and associated secretory vesicles are proposed to be involved in microneme and rhoptry spatial organisation within the conoid, but optical resolution has been insufficient to confirm this hypothesis. Recently, a number of molecules involved in rhoptry discharge have been identified, including orthologues of *Plasmodium* non-discharge rosette complexes, as well as rhoptry apical surface proteins (Suarez et al., 2019) and calcium-sensing ferlins (Coleman et al., 2018). While these molecules may hold the key to signalling, the key mechanisms of rhoptry exocytosis, whereby RONs and ROPs cross both the parasite and the host PMs, remain mysterious. The discharge of rhoptry and microneme proteins occur at the apical end presumably by fusion with the PM overlying the conoid (Bannister et al., 2003; Scholtyseck and Mehlhorn, 1970), but this is not well understood and there is uncertainty as to the machinery involved in fusion prior to release. A recent study using cryo-electron tomography of *Toxoplasma gondii* tachyzoites showed rhoptries connected to an apical vesicle underlying the plasma membrane of the conoid and a rosette of non-discharge proteins embedded in the parasite plasma membrane, which appears to represent machinery at least in part required for fusion and delivery of rhoptry contents into the host cytosol (Aquilini et al., 2021).

Rhoptries have been visualised within the conoid in many studies (Dos Santos Pacheco et al., 2020), but there has been some uncertainty over the precise location of micronemes either within the conoid (Dubois and Soldati-Favre, 2019) or at the base of the conoid when it is protruded (Paredes-Santos et al., 2012). Here, we used high resolution cellular electron tomography to reconstruct the apical complex of *Eimeria tenella* sporozoites. Data from serial electron microscopy tomography and serial block-face scanning electron microscopy (SBF-SEM) were combined to produce 3D models showing how secretory organelles are arranged for exocytosis. Our data provide evidence for fusion of the rhoptry membrane with the parasite membrane overlying the conoid and a pore in the parasitophorous vacuole membrane that may be important in the delivery of rhoptry contents to the host cytosol early in infection.

## Results

### Cellular electron tomography of the apical complex reveals a highly ordered gateway for secretion

To investigate the detailed ultrastructure of the conoid and how the secretory organelles are organised in the apical complex, serial section electron tomography was performed on freshly hatched extracelluar sporozoites and invaded sporozoites (N=17, serial dual axis serial tomograms). These tomograms encompassed the entire apical end covering ~2.5 μm^3^. Segmentation of the conoid and associated structures were carried out for each serial tomogram (e.g. Movie 1 and selected slices through a serial section tomogram images in Fig. 1C). Due to the complexity of the overall structure, we divided the apical complex in two parts: the structures/organelles that surround the conoid barrel are referred to as the ‘outer’ conoid components (Fig 1A) and the structures/organelles contained within the barrel of the conoid are called ‘inner’ conoid components (Fig 1B). The outer apical complex includes an electron-opaque apical polar ring (APR) (Fig 1A; orange). Our work confirms that there are 24 evenly spaced subpellicular microtubules (Fig 1A; green) radiating from APR and extending towards the posterior of the cell, as identified in all 17 tomograms. Two ring-shaped structures distal to the conoid - known as pre-conoidal ring-1 (PCR-1) (Fig 1A, light blue) and pre-conoidal ring 2 (PCR-2) (Fig 1A, red) - complete the outer conoid components. The outer apical complex is enclosed by the plasmalemma and partially enclosed by the inner membrane complex (IMC), which is interrupted apically to form a circular apical opening (not shown in the tomograms). This is located distal to the apical polar ring (APR).

**Figure 1:**
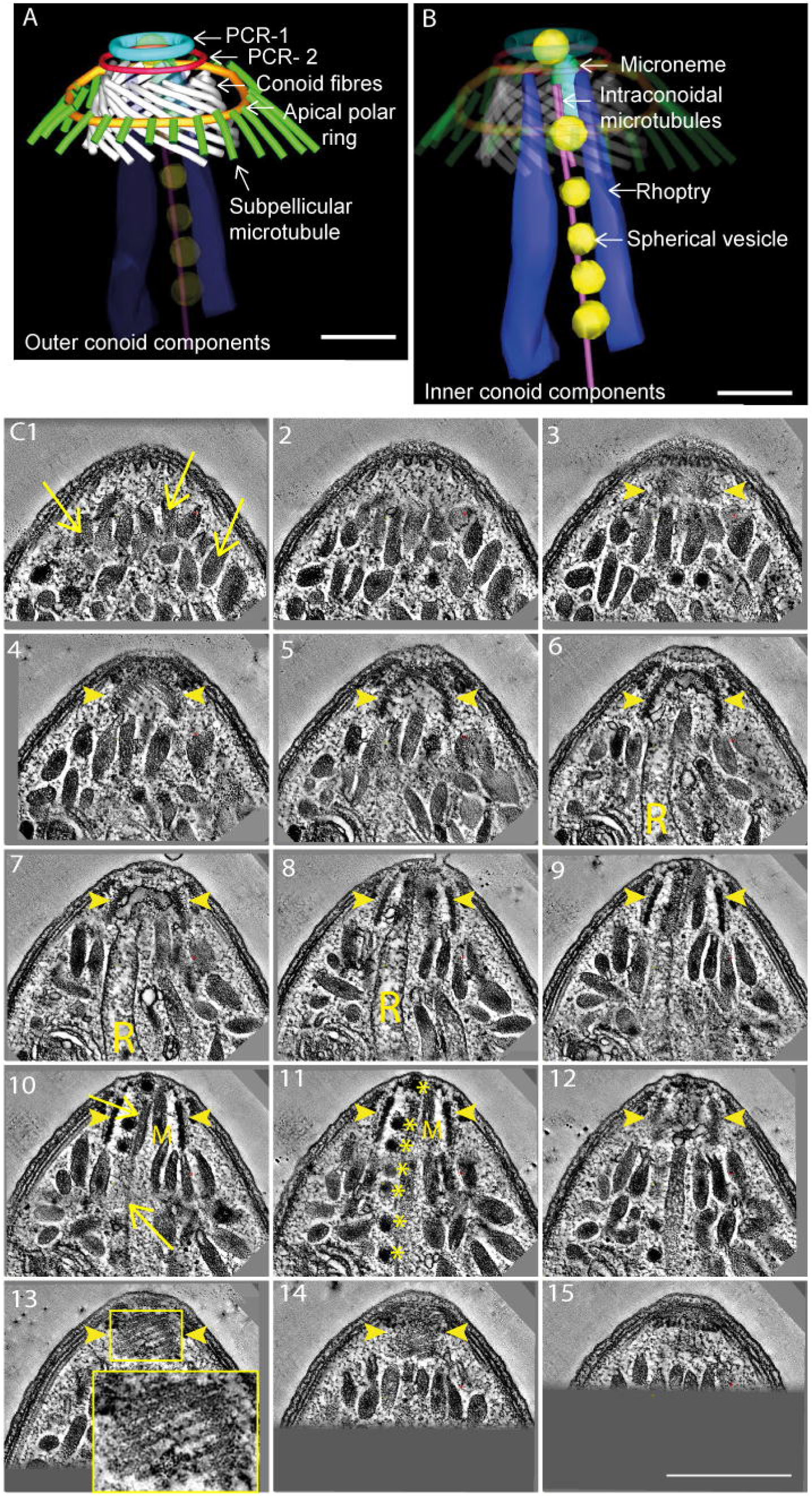
Components of the apical complex in sporozoites of *E. tenella*. The apical complex is divided into outer conoid (A) and inner conoid (B) components. A and B: Segmentation from tomogram. Outer conoid components; Conoid fibres (white); 2 pre-conoidal rings (PCR 1 and 2) (light blue and red), apical polar ring (gold) in association with sub-pellicular microtubules (green); B: Inner conoid components, rhoptry (dark blue), microneme (light green), secretory vesicle (yellow), intra-conoidal microtubule pair (pink); Scale bars - 200nm. C: Series of 15 tomographic slices through a representative tomogram, conoid – yellow arrowheads, micronemes – arrows throughout the series, microneme within the conoid (10,11), Rhoptry (7, 8), long and short intra-conoidal microtubules (10), spherical vesicle asterisks (11), Conoid fibres and inset (13). Scale bar - 500nm.

The inner apical complex components comprise a pair of centrally located intra-conoidal microtubules (Fig 1B; pink). We discovered that one of these microtubules starts and terminates in line with the height of the conoid, whilst the other extends into the cell interior past the base of the conoid and beyond the region covered in our tomograms, which has not been reported previously (Fig 1B; pink, 1C:10 – arrows, 1C:11). Spherical vesicles of varying number are closely associated with the two intra-conoidal microtubules and appear as a row of electron-opaque spheres leading posteriorly from each conoid (Fig 1B – yellow; 1C: 11 - asterisks). Spherical vesicles have been identified in this configuration in numerous studies, but the composition of these are unknown (Paredes-Santos et al., 2012). There has been some uncertainty over the presence or absence of microneme trafficking into the conoid barrel in *Toxoplasma gondii* (Dubois and Soldati-Favre, 2019), but our tomograms clearly show both rhoptries (Fig 1B, dark blue; 1C: 7,8 - “R”) and micronemes (Fig 1B, light green; 1C: 10 - “M”) located within the conoid area of all 17 tomograms (see below, movie 1, Supple. Fig 2).

When viewed as a three-dimensional reconstruction, the conoid appears as an open truncated cone formed from closely apposed helical fibres which follow a left-handed helical path towards the apical end of the parasite (Fig 1A; white). Quantification of the size of the conoid and the number of tubulin-containing fibres reveals a variation in fibre number between individual parasites, but in all tomograms there were always 24 evenly spaced subpellicular microtubules connected to the apical polar ring that surrounds the conoid. Negatively stained whole-mount cytoskeletons of *T. gondii* estimates that the conoid is composed of ~14 helically arranged tubulin-containing fibers (Hu et al., 2002), but high resolution tomography of the conoid had not been carried out to confirm this. Our high-resolution tomograms show that there are between 13 and 16 fibres in the conoid, and there can be variation in fibre number even in genetically identical sporozoites within the same sporocyst (Shirley and Harvey, 1996) (Supple. Fig. 1A). The width of each conoid fibre was constant at 22 ± 2 nm and the spacing between each conoid fibre also remained constant at 33 nm ± 4 nm irrespective of conoid fiber number (supple. Fig 1B). This fixed arrangement of fibre width and distance between each fibre means that a conoid with more fibres must have a larger diameter (Supple. Fig 1C). This was confirmed by measurements of the conoid diameter at the base (mean = 340 nm, +/−20 nm SD) and correlated with fibre number, which revealed a positive correlation (coefficient = 0.78, p = 0.014, Pearson correlation).

Further quantitative analysis of the relationship between the increasing conoid diameter and the size of other apical structures was carried out. There was no correlation between conoid height (mean = 193 nm, +/− 35 nm SD) and fibre number (coefficient = 0.57, p=0.113, Pearson correlation), suggesting that differences in fibre number are not linked the length of the conoid fibres (Supple. Fig 1C). The conoid diameter did correlate to the diameter of the pre-conoidal rings. PCR-1 diameter was 169.3 nm (+/− 9.6 nm) and PCR-2 diameter was 221.5 nm (+/− 14.5 nm) and the increase in diameter of both rings was correlated with increasing conoid base diameter (PCR-1; Pearson correlation, r = 0.97, p<0.001; PCR-2; Pearson correlation, r = 0.74, p = 0.037). Finally, APR diameter was 393nm (+/− 19 nm) and there was also a positive correlation between APR diameter and conoid base diameter (= 0.76; p = 0.017, Pearson correlation). Presumably, this would be important in order to maintain the distance between the APR and conoid (Supple. Fig 1D). In summary, our measurements reveal there are differences in conoid fibre number resulting in dependency relationships between the size of the conoid and outer conoid components in order to maintain the integrity of the overall structure despite size heterogeneity between individual parasites.

### Micronemes and rhoptries are organised along the intra-conoidal microtubules and closely associated with the plasma membrane overlying the conoid

The serial tomograms we generated encompassed the full area of the conoid in enough resolution to allow us to establish the precise location and number of each type of secretory organelle. Although microneme proteins are known to be secreted at the apical pole, the precise location and mechanism of trafficking of microneme proteins into the apical complex is uncertain and a detailed quantitative analysis had not been carried out previously. In our dataset set we discovered 1-2 rhoptries, 1-5 micronemes and 1-3 spherical vesicles within the barrel of the conoid in all tomograms (Fig 2A-D, Fig 1L, M). Supple. Fig 2 shows examples of 5 freshly excysted sporozoites with differing number of micronemes within the conoid area and entering the conoid area. In addition, rhoptries, micronemes and spherical vesicles were always closely associated with intra-conoidal microtubules, which are located centrally within the barrel of the conoid (Fig 1B, 2A, Movie 1). To quantify the spatial relationships between the intra-conoidal microtubule pair and micronemes, rhoptries and spherical vesicles, we measured the distances between each organelle and the intra-conoidal microtubules, and between each organelle and the conoid fibres (Fig 2C - G). One of the organelles were chosen in each tomogram and measurements taken at roughly the same position within the conoid to take account of the cone shape of the conoid. These measurements reveal that micronemes, rhoptries and spherical vesicles are always located significantly closer to the intra-conoidal microtubule pair within the central barrel of the conoid than to the fibres of the conoid barrel (Fig 2E-G), suggesting a potential role for the intra-conoidal microtubules in organising the secretory organelles within the conoid in readiness for secretion. The plasma membrane overlying the conoid was checked for evidence of fusion events or close apposition of microneme, rhoptry or spherical vesicle membranes. All rhoptries within the conoid were in close apposition with the plasma membrane. Spherical vesicles were organised in single file along the intra-conoidal microtubules and there were always 1 or 2 micronemes and at least 1 spherical vesicle located in close apposition with the plasma membrane overlying the conoid, as well as being closely associated with the intra-conoidal microtubules (Fig 2H-O). This suggests there is a sequential trafficking of micronemes and spherical vesicles along the intra-conoidal microtubules to the plasma membrane overlying the conoid, in preparation for secretion.

**Figure 2:**
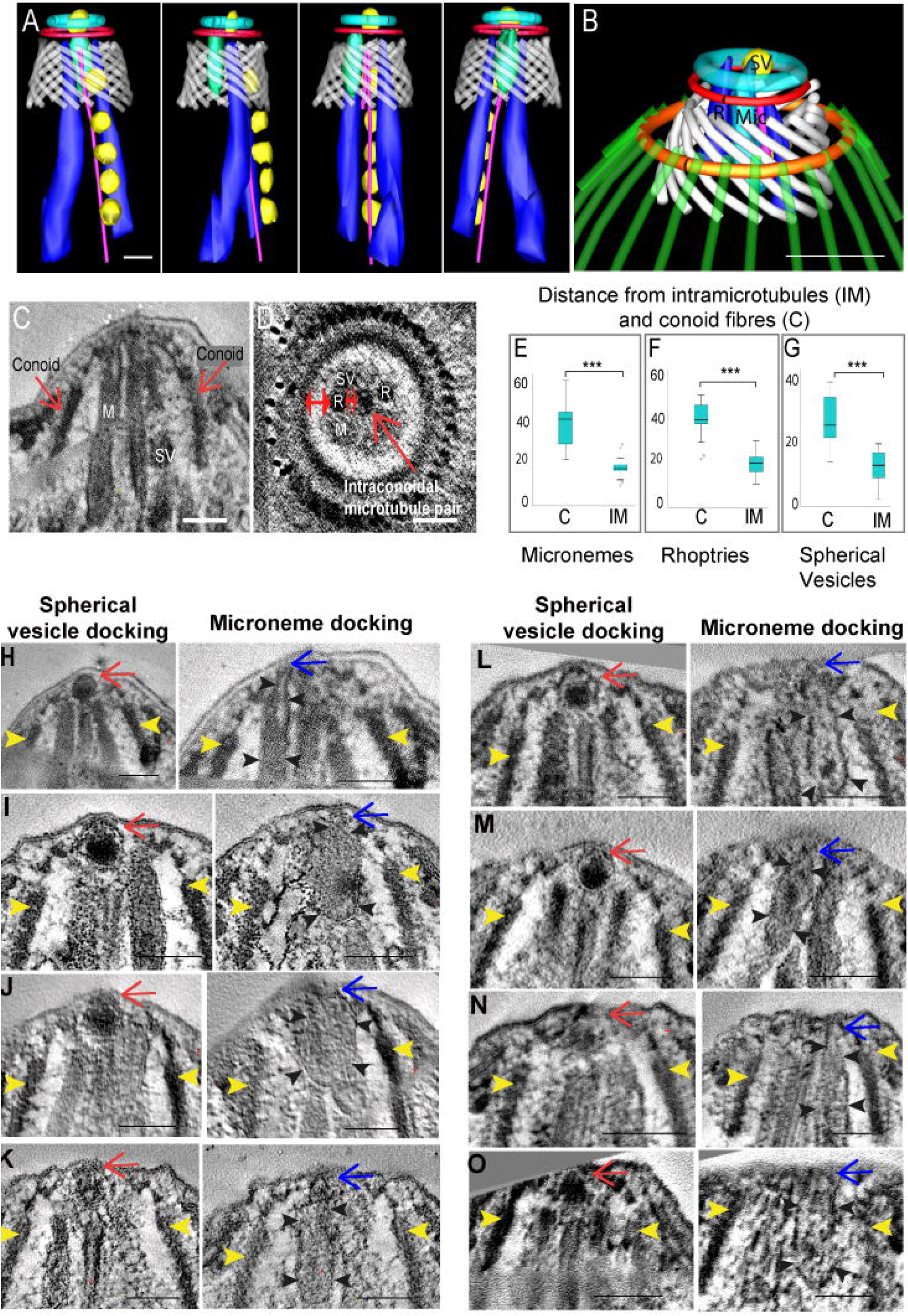
Secretory organelles and their association with the intra-conoidal microtubules and overlying the plasma membrane. A: Series of rotational views of a segmentation created from one serial tomogram illustrating the spatial grouping and alignment of microneme (light blue), rhoptries (dark blue) and spherical vesicles (yellow) with the intra-conoidal microtubule pair (pink); B: Segmentation of a serial tomogram illustrating the relative positioning of the microneme (light green), rhoptry (dark blue) and spherical vesicles (yellow) with the apical end of the parasite; C and D: slices taken from different tomograms (C) longitudinal view and (D) cross section view illustrating how measurements were taken showing the distances from a microneme (M), rhoptry (R) and spherical vesicle (SV) to either the intra-conoidal microtubules or conoid. Double-headed red arrows show where the measurements were taken, single arrow in C for location of the conoid. Single arrow in D the location of intra-conoidal microtubules; E-G: Micronemes, spherical vesicles and rhoptries were significantly closer to the intra-conoidal microtubules (IM) than to the conoid (C) (t-test, p < 0.0001) (N= 17); H - I: Longitudinal slice views from 8 tomograms illustrating a spherical vesicle (red arrows), microneme (blue arrows and black arrowheads) in close association with the plasma membrane overlying the conoid in each tomogram. Yellow arrowheads show the outer edge of the conoid in each example. Scale bars – 100nm.

Whilst fusion of microneme or spherical vesicle membranes with the plasma membrane was not observed in freshly excysted sporozoites, our dataset included two serial section dual-axis tomograms of sporozoites situated within a parasitophorous vacuole (PV) after sporozoite invasion of Madin-Darby bovine kidney (MDBK) cells. In one of these invaded sporozoite tomograms fixed at 30 min post-infection, a rhoptry was identified with an electron-lucent interior instead of the usual electron dense rhoptry organisation (Fig 3A and inset, B and inset; Movie 2). The electron-lucent rhoptry (ELR) differed in shape from the electron dense rhoptries (R) in the same and other tomograms, having a broader base and flask-shaped (Fig 3C). Intriguingly, the electron-lucent rhoptry appeared to be continuous with the parasite plasma membrane as if the rhoptry had fused with the parasite plasma membrane overlying the conoid, to release its contents (Fig 3B and inset). At the point of fusion there was a hole in the PV membrane (PVM) appearing to create a channel passing through both the parasite plasma membrane and the PVM, connecting the electron-lucent rhoptry directly with host cell cytoplasm (Fig 3B and inset, C, D, Movie 2).

**Figure 3:**
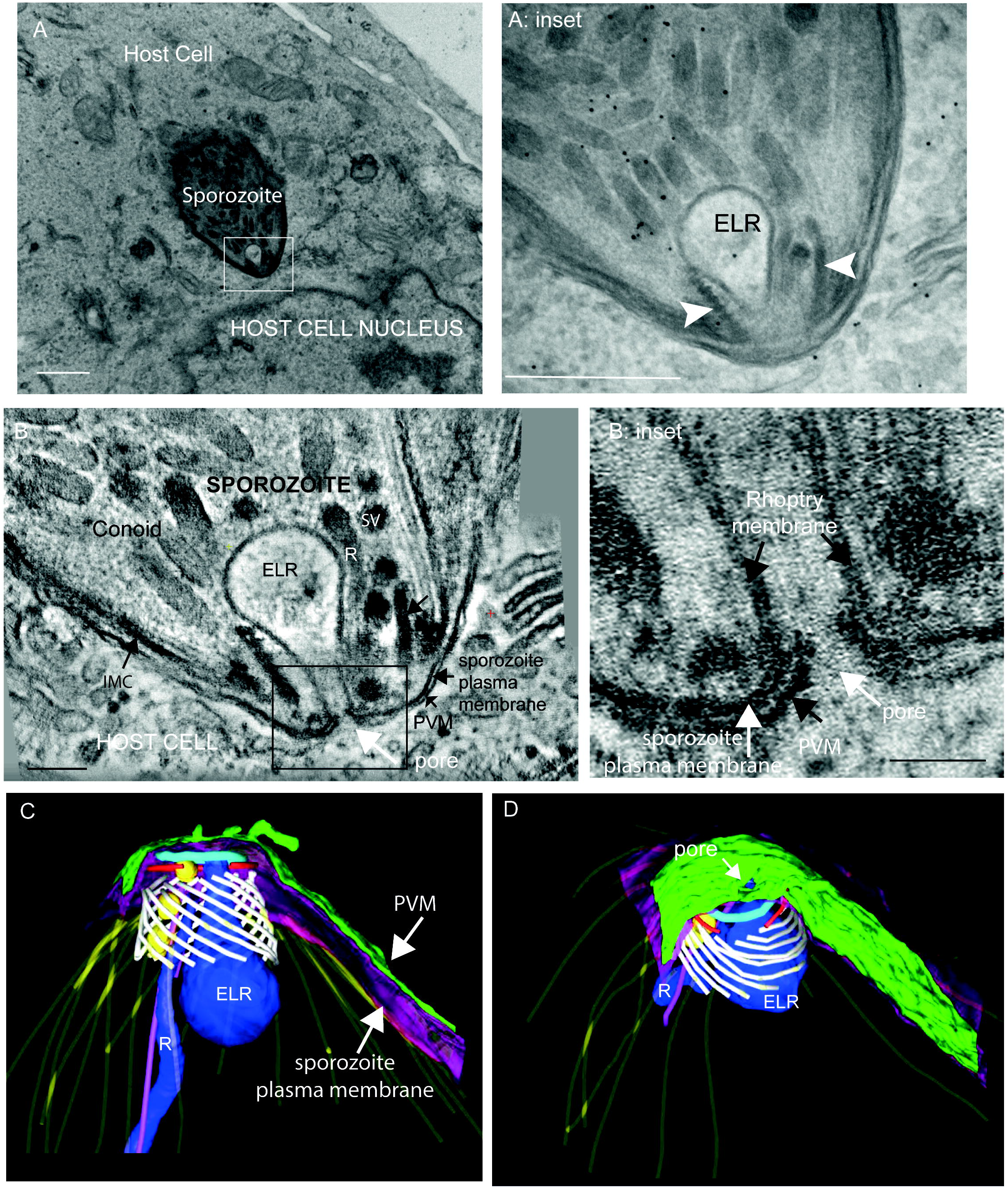
Characterisation of an electron lucent rhoptry in an intracellular sporozoite. A: Slice from a tomogram illustrating an intracellular sporozoite with an electron lucent rhoptry (box and A: inset) close to a host cell nucleus (30 min post-infection sample) scale bar 1μm; A: inset: Higher magnification of A illustrating the electron lucent rhoptry (ELR) at the conoid (white arrowheads point out the boundaries of the conoid). Scale bar 500nm; B and B: inset: Slice from a tomogram showing the electron lucent ‘empty’ rhoptry (ELR) which appeared to be continuous with the parasite plasma membrane and attached at its apex to a hole in the parasitophorous membrane creating a channel passing through both the parasite plasma membrane and the PVM connecting the ‘empty rhoptry’ directly with host cell cytoplasm B – scale bar 100nm; B inset scale bar 50nm; C and D: Segmentation of a serial tomogram outlined in A and B to illustrate the three dimensional organisation and relative positioning of the electron lucent rhoptry (ELR – dark blue), electron dense rhoptry (R – dark blue), conoid fibres (white), spherical vesicles (yellow), parasite plasma membrane (purple), parasitophorous vacuole membrane (green), PCR-1 and 2 (light blue and red), subpellicular microtubules (yellow), intra-conoidal microtubules (pink).

Overall, our extensive tomography analysis of both freshly excysted and invaded sporozoites revealed that different types of secretory organelles converge within the conoid and are closely associated with the intra-conoidalal microtubules, forming what appears to be a highly ordered secretory gateway at the apical end of the cell.

### Whole cell reconstructions of individual sporozoites reveals the organisation and location of major sporozoite organelles

Our tomography data revealed that only a limited number of individual secretory organelles are located within the conoid area, yet many studies show the presence of hundreds of micronemes within the apical end of the parasite outside the conoid area. To better understand the broad localisation and number of secretory organelles, freshly excysted sporozoites were prepared for serial block face-scanning electron microscopy (SBF-SEM), which allows automated collection of datasets containing hundreds of sequential serial sections. A total of 25 whole individual sporozoite cells found in aligned SBF-SEM series were reconstructed and analysed for organelle number and three dimensional organisation. A single slice of a freshly excysted sporozoite illustrates that most major organelles were visible by SBF-SEM (Fig 4A, B, C; modelled in 4F). Movie 3 illustrates a portion of the SBF-SEM dataset showing 25 slices containing a sporozoite. Organelle volume and number were determined by manual segmentation of each organelle, and the relative abundance and location of all identifiable organelles (including micronemes and rhoptries) were analysed (Fig 4F; Suppl. Fig 3). Due to the number and close packing of micronemes located mainly at the apical end of the sporozoite, it was not possible to accurately count them. Instead, micronemes were segmented using a combination of pixel-density thresholding and manual area selection in three whole cell volumes (Fig 4D). The total mean whole cell volume of individual sporozoites was 61.27 μm^3^ (Supple. Fig 3A) with micronemes comprising 5% of whole cell volume and rhoptries <0.5% of whole cell volume (Fig 4G). This quantitative analysis revealed that refractile bodies make up 36% of whole cell volume (2 per cell in freshly excysted sporozoites), with the nucleus (1 per cell) and amylopectin granules (~196/cell) each comprising 4%, mitochondria (~14/cell) 2% and acidocalcisomes (~13/cell) 1% of whole cell volume (Supple. Fig 3 for full quantification). The majority of micronemes were densely packed at the apical end of the sporozoite (Fig 3D). Rhoptries were observed as club-shaped structures with a rounded ‘bulb’ region and an elongated ‘neck’ region. They were also mostly found towards the sporozoite apical end, although some rhoptries were present in the central and posterior parts of the cell (Fig 4E). Unfortunately, the spherical vesicles observed by tomography (in association with the intra-conoidal microtubules, e.g Fig 1B) were not clearly identified in the reconstructed whole cell SBF-SEM data. By combining findings from tomography and SBF-SEM, we conclude that *E. tenella* sporozoites contain an abundance of micronemes and rhoptries, and only a small number of these are seen within the conoid where secretion occurs, suggesting that there is significant directed movement of these secretory organelles converging at the conoid.

**Figure 4:**
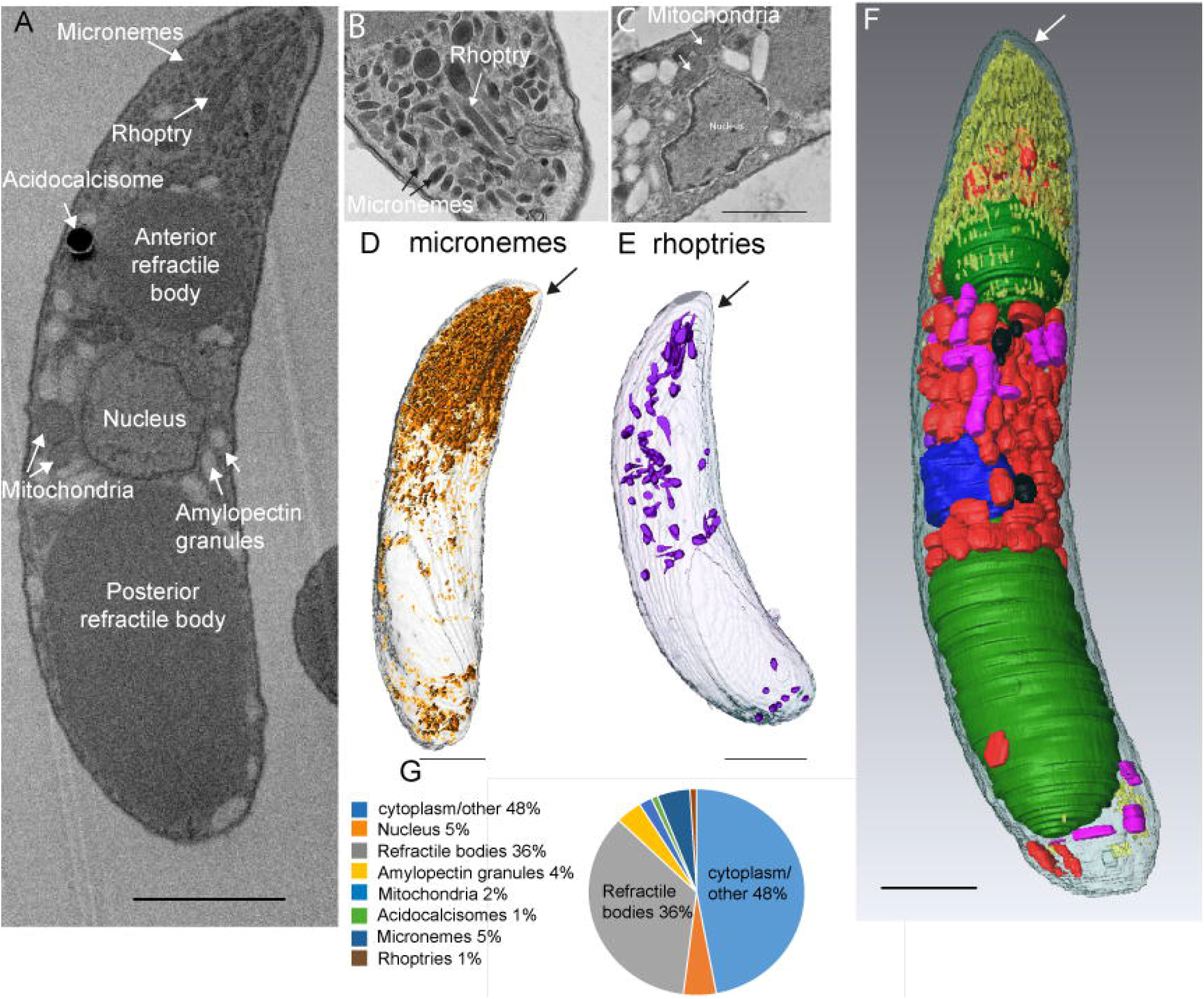
SBF-SEM quantification of micronemes, rhoptries and other major organelles in freshly excysted sporozoites. A: Longitudinal section from an SBF-SEM dataset illustrating the major organelles. B and C: Additional slices from SBF-SEM datasets to illustrate identification of major organelles. D and E: Quantification and location of microneme and rhoptry organelles in SBF-SEM whole cell reconstructions. F. Combined model illustrating the positioning of all the major organelles: micronemes – yellow, refractile bodies – green, amylopectin granules – red, acidocalcisomes – black, rhoptries – purple, nucleus – blue - Scale bars – 1μm; G. Relative volumes of major organelles in freshly excysted sporozoites.

## Discussion

Secretion of microneme and rhoptry contents from the apical end of apicomplexan zoites is well documented and hundreds of micronemes have been visualised closely packed at the apical end of many different apicomplexan parasites by classic transmission electron microscopy (TEM) thin sections (For review see Dubois and Soldati-Favre, 2019). Exactly how micronemes are trafficked for secretion to this small portion of plasma membrane is not well understood. In *T. gondii*, focussed ion beam scanning electron microscopy (FIB-SEM) images did not find micronemes within the conoid area and it was proposed that microneme secretion could occur adjacent to the conoid, close to APR and the subpellicular microtubules, where a membrane space could open up as the conoid protrudes (Paredes-Santos et al., 2012), rather than secretion occurring at the plasma membrane directly overlying the conoid. However, further studies using the same technique did observe a microneme inside the conoid area. Although the intra-conoidal microtubules could not be resolved by FIB-SEM (Dubois and Soldati-Favre, 2019), they have been proposed to be involved in secretory cargo trafficking (Hu et al., 2006b). Our experimental data using high resolution cellular electron tomography in *E. tenella* provide experimental evidence for the role of the intra-conoidal microtubules in the trafficking of micronemes through the conoid to the plasma membrane overlying the conoid, suggesting an orderly system for transporting these organelles to the membrane in preparation for docking, fusion and release of contents. We also show here that rhoptries and a set of spherical vesicles (of yet unknown composition) are also closely associated with the intra-conoidal microtubules. This suggests that the intra-conoidal microtubules are a major organiser of orderly trafficking within the conoid area. In addition, our tomograms show that at least one intra-conoidal microtubule may be involved in trafficking of organelles into the conoid space, because this microtubule extends posteriorly into the cytoplasm (beyond the region covered by our tomograms), where it is ideally positioned to interact with secretory organelles and ‘guide’ them to the conoid for discharge.

Careful analysis of tomograms did not find direct evidence of fusion of microneme or spherical vesicle membranes with the plasma membrane overlying the conoid, despite their close proximity to the plasma membrane in nearly all tomograms. This might be due to the fixation method used in this study or could indicate that the process is very fast and therefore extremely difficult to capture in still images, but certainly the signalling cascade for membrane fusion exists in apicomplexan organisms (for review see (Dubois and Soldati-Favre, 2019). Our observation of an apparent fusion of a rhoptry to the parasite plasma membrane overlying the conoid and a small pore in the parasitophorous vacuole membrane surrounding an invaded sporozoite has significant implications in re-shaping what is understood about the parasitophorous vacuole. The current view of the apicomplexan parasitophorous vacuole membrane is that this is sealed, requiring insertion of parasite-derived pores for the transport of molecules between the host cell cytosol and the parasitophorous intra-vacuolar space, and in particular for the trafficking of rhoptry proteins into the host cytosol (Gold et al., 2015; Marino et al., 2018). However, our data as well as a rare thin section electron microscopy of an invaded *T. gondii* tachyzoite show a direct connection or pore linking an electron-lucent rhoptry and the host cell cytosol (Nichols et al., 1983). Recently, a mechanism proposed to be involved upstream of rhoptry fusion was discovered in *T. gondii* tachyzoites using cryo-electron tomography. The authors showed that rhoptries connected with an apical vesicle underlying the plasma membrane of the conoid and the position of the apical vesicle coincided with a rosette of non-discharge proteins embedded in the parasite plasma membrane. Direct fusion and a pore were not observed in this study as we show in this work, but the formation of a rosette containing non-discharge proteins maybe a pre-requisite for fusion of membranes and release of rhoptry proteins (Aquilini et al., 2021). It is probable that rhoptry fusion is a highly dynamic event that takes place only at specific stages of intracellular infection, which might explain why they have been visualised only on rare occasions. In addition the pore is small (~40nm), which would make it even more difficult to obtain clear images by thin section transmission electron microscopy.

Our detailed measurements and quantification of the conoid showed a highly ordered organisation of conoid fibres in a left-handed helical organisation with fixed spacing between the fibres. This is the first detailed high resolution three dimensional reconstruction of a coccidian conoid and interestingly we have shown there is heterogeneity in fibre number between individual sporozoites even at different developmental stages. This difference in fibre number influences the overall size of the conoid and directly correlates with the associated diameters of the two conoidal rings and the APR, presumably to ensure overall structural integrity to allow for conoid mobility, so it can be recessed and flushed with the apical polar ring or extruded beyond the apical polar ring (Monteiro et al., 2001). Intriguingly in all parasites there were always 24 subpellicular microtubules on the APR, which presumably places a physical constraint for the minimum and maximum possible diameter of the APR and its associated structures and thus, dictates the minimum and maximum overall dimensions of the apical complex at least in the sporozoite stage of the parasite.

Here we show that combining high resolution cellular electron tomography and lower resolution SBF-SEM data is a powerful way of investigating specific areas of cells with a whole cell view. These datasets reveal a highly organised gateway for trafficking of secretory organelles to the conoid area of the apical complex. Further work will be required to understand the role of rhoptry fusion and pore formation within the PV and how individual micronemes are trafficked into the conoid area from such a large cluster underlying the conoid area.

## Acknowledgements

This research was supported by a joint PhD studentship between Oxford Brookes University, Oxford, UK and The Royal Vet College (RVC), University of London to Alana Burrell. VMH was funded by BBSRC grant BB/L00299X/1 and by a research fellowship from the RVC. We would like to thank staff at the Oxford Brookes Centre for Bioimaging for technical assistance and advice during collection of datasets and to staff at the Biological Services Unit at the RVC for their assistance in the care of the animals. We also thank Ryuji Yanase and Heloise Gabriel both Oxford Brookes University for assistance in data analysis.

## Figure legends

**Supplemental figure 1: Conoid fibre number variation in sporozoites from pre-excystation sporocysts, freshly excysted and intracellular.** A: Segmentation of sporozoite conoids from pre-excystation sporozoites within sporocysts, freshly excysted and intracellular sporozoites. The numbers next to sporocysts indicates matching sporozoites within a sporocyst; B: segmentation of a conoid (white) illustrating how measurements were taken for conoid fibre width (1) and conoid fibre to fibre spacing (2); C: segmentation of sporozoite conoid area illustrating how measurements of conoid diameter and height were taken; D: segmentation of a sporozoite conoid area illustrating the measurement criteria of conoid to apical polar ring (arrow). Index: Conoid fibres (white); 2 pre-conoidal rings (PCR 1 and 2) (light blue and red), apical polar ring (gold) in association with sub-pellicular microtubules (green).

**Supplemental figure 2:** A selected tomogram slices from 5 serial tomograms to illustrate the number of micronemes (mic) and rhoptries in the conoid area. All micronemes that were either partially or fully within the conoid were included. Conoid is highlighted with yellow arrowheads in all examples. Tomogram A containing 2 rhoptries and 5 micronemes within the conoid. Mic 1 and 3 are closest to the plasma membrane underlying the conoid. Mic 2, 4 and 5 have partially entered the conoid area; Tomogram B, slices from a tomogram containing 2 rhoptries and 3 micronemes. Mic 1 is closest to the plasma membrane; Tomogram C, slices from a tomogram with 2 rhoptries and 2 micronemes; Tomogram D, slices from a tomogram containing 2 rhopries and 2 micronemes. Mic 1 is closest to the plasma membrane; Tomogram E, slices from a tomogram containing 2 rhoptries and 2 micronemes.

**Supplemental figure 3: Volume and numbers of major organelles from whole cell reconstructions of freshly excysted sporozoites by SBF-SEM.** Analyses was calculated from segmented SBF-SEM data for each organelle in freshly excysted sporozoites. For each organelle the mean volume of an individual organelle is included, SD = standard deviation, COV = co-efficient of variation, range of volumes of a particular organelle or cell volume. The number of organelles per cell is included for amylopectin granules, acidocalcisomes and mitochondria. AP = Apical end of the parasite. Scale bar 1μm.

**Movie 1:** Serial section cellular electron tomogram containing the apical complex of a freshly excysted sporozoite and segmentation of the dataset to illustrate the three dimensional model. Conoid fibres (white); 2 pre-conoidal rings (PCR 1 and 2) (light blue and red), apical polar ring (gold) in association with sub-pellicular microtubules (green); rhoptry X 1 within the conoid (dark blue), micronemes X 2 modelled within the conoid (light green), secretory vesicles (yellow), intra-conoidal microtubule pair (pink);

**Movie 2:** Serial section cellular electron tomogram of an infected MDKB cell 30 mins post-infection containing an electron lucent rhoptry and pore. Movie illustrates the tomogram data followed by the segmentation. Colour scheme as per Movie 1 figure legend.

**Movie 3:** A total of 25 sequential slices (~100nm thick) through an SBF-SEM dataset to illustrate a whole freshly excysted sporozoite used for quantitative analysis of organelles in Fig 4 and Supple. Fig 3.

## Methods

### Infection of chickens and sporozoite purification

Three-week-old Lohmann chickens (purchased from APHA Weybridge) kept under specific pathogen free conditions were orally infected with 4,000 sporulated *E. tenella* Wisconsin strain oocysts (Shirley, 1995). Oocysts were harvested at 7 days post-infection and excystation and sporozoite purification performed as previously described (Pastor-Fernández et al., 2020).

### In vitro *E. tenella* infections

The NBL-1 line of MDBK cells (ECACC-Sigma-Aldrich, Salisbury, UK) were prepared as previously described (Marugan-Hernandez et al; 2020). One millilitre of cell-culture medium containing 0.35 x 10^6^ MDBK cells was added to wells of a 24 well culture plate. Cells were left to settle for up to 3 hr at 38 ⁰C, 5% CO_2_, prior to infection with sporozoites. Freshly-purified sporozoites were pelleted by centrifugation at ~ 600 RCF for 10 min and re-suspended in cell-culture medium at a concentration of 3.5 million sporozoites per ml. One millilitre of sporozoite suspension was added to each MDBK-cell containing well.

### Preparation of freshly excysted sporozoites for electron microscopy

Freshly purified sporozoites (~10-50 million) were suspended in 1 ml of primary fix (2% freshly-prepared formaldehyde solution (Sigma-Aldrich), 2.5% electron microscopy grade glutaraldehyde (TAAB) and 0.1 M sodium cacodylate buffer (TAAB) in double distilled (dd H_2_O). Sporozoites were left in primary fixative for two hours at 4°C. Fixed sporozoites were washed five times in 0.1 M cacodylate buffer pH 7.4 for 10 min. Sporozoites were pelleted by centrifugation and incubated in 2% osmium tetroxide (TAAB) in 0.1 M cacodylate buffer for 60 min at 4°C. For uranyl acetate staining, sporozoites were added to 1 ml molten 3% agarose (2-Hydroxyethyl agarose – Sigma-Aldrich, dissolved in ddH_2_O), centrifuged and incubated at 4°C for 5 min. Approximately 1 mm³ blocks were cut from the part of the agarose containing sporozoites and incubated in freshly-filtered 2% aqueous uranyl acetate (Agar scientific) in the dark at 4°C overnight. Sporozoites were washed in ddH_2_0 and then dehydrated by a series of 20 min incubations in acetone/ddH_2_0 solutions. Samples were incubated for 2 hours in 25% epoxy resin (TAAB 812 resin premix kit) in acetone; overnight in 50% resin in acetone; 6 hrs in 75% resin in acetone; overnight in 100% resin, and finally, two changes of 100% resin for 2 hours each. Polymerisation was achieved by incubation at 60°C for 24 h.

### Preparation of host cells infected with sporozoites for electron microscopy

Electron microscopy grade glutaraldehyde was added to the infected MDBK cells for 15 min then primary fixation carried out as above.

### Electron tomography and measurements

Transmission electron tomography was performed using one of several transmission electron microscopes: H-7560 (Hitachi™), Spirit (Tecnai™, FEI™/Thermo Fisher Scientific™) or Talos™ (FEI™/Thermo Fisher Scientific™). Sections were cut at 150 nm thickness for 120 kilovolts (kV) (Hitachi™ H-7560 or Tecnai™ Spirit electron microscopes) and at 150 nm-200 nm thickness for 200 kV (FEI™ Talos™ electron microscope). Tomogram tilt series generated using the Hitachi™ H-7560 electron microscope were taken from −60° to −60° with intervals of −1°. For tomography data acquisition using the Tecnai™ Spirit electron microscope, automated centring, focus adjustment, tilt setting, and image capture were performed using Xplore*3D*™ software by FEI™. For tomography data acquisition using the FEI™ Talos™ electron microscope, automated centring, focus adjustment, tilt setting, and image capture were performed using DigitalMicrograph™ (Gatan™) with SerialEM™ plug-in (Mastronarde, October 2005). Regardless of the microscope used, dual axis tomograms were collected by rotating the grid by 90° and repeating the tilt series image collection. Tomogram image data series processing, segmentation of the tomograms (modelling) to produce three dimensional reconstructions and all measurements from the tomograms were also carried out using using IMOD™ software (Kremer et al., 1996) (University of Colorado, Boulder). All measurements were carried out in IMOD. Measurement were also carried out using the measurement features in IMOD.

### Serial block face scanning electron microscopy (SBF-SEM) image acquisition

SBF-SEM data was collected using a Zeiss™ Merlin scanning electron microscope with Gatan™ 3View™ automated sectioning and image capture system. Samples were trimmed to ~1 mm^3^ and mounted on 3view sample pins using an epoxy conductive adhesive from Circuitworks™. After insertion of the mounted block, the intra-microscope diamond-knife was advanced towards the block-face by 200 nm cutting-strokes until sectioning of the block face was observed. The block-face was then imaged using a scanning electron beam with 3-5 kV accelerating voltage. Electron signal was detected using a back-scatter electron detector (OnPoint, Gatan) and nitrogen gas was injected to raise chamber pressure to 30pa. SBF-SEM data was processed using IMOD™ software, run through Cygwin™ command line interface. Segmentation was also carried out using IMOD™ software. (University of Colorado, Boulder).

### Statistical analysis

Statistical analyses were performed using IBM™ SPSS™ version 25 software. A t-test was used to test for an association between a continuous variable and a binary categorical variable where there was normal distribution for both groups. If testing for an association between a continuous variable and a binary categorical variable where one or both groups did not show normal distribution, the Mann-Whitney U test was used. If dealing with a continuous variable and a categorical variable with more than two groups, where the continuous data was normally distributed for all the groups, One-Way ANOVA was used followed by a post-hoc Turkey HSD test. If there was a continuous variable and a categorical variable with more than two groups, where data within at least one of the groups was not normally distributed, then the Kruskal-Wallis test was used. The Chi-squared test was used when assessing for a statistically significant association between two categorical variables. See individual figures and legends for specific tests for each experiment.

## Notes

### Competing Interest Statement

The authors have declared no competing interest.

