## supplemental figure for "Cellular electron tomography of the apical complex in the apicomplexan parasite *Eimeria tenella* shows a highly organised gateway for regulated secretion"

### Supplemental figure 3

#### Cell Volume

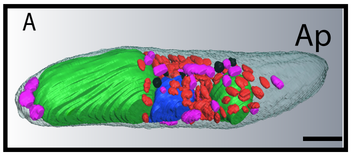


Mean = 61.27 µm3
SD = 10.9 µm3
COV = 17.7%
Range = 36.5 µm3- 72.4 µm3

#### 1 x Nucleus

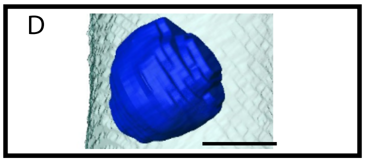


Mean = 2.3µm3
SD = 0.4µm3
COV = 17.2%
Range = 1.58µm3 - 2.83µm3

#### 2 x Refractile Bodies

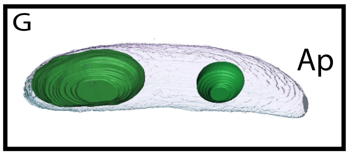


Mean = 22.1µm3–
SD = 5.45µm3
COV = 24.7%
Range = 10.51µm3 - 29.2µm3

#### Mitochondria

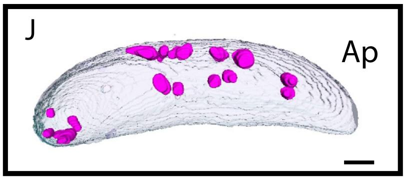


**Volume:**
Mean = 1.14µm3
SD = 0.31µm3
COV = 27.1%
Range = 0.47µm3 - 1.82µm3

**Number:**
Mean = 14
SD = 5.4
COV = 38.7%
Range = 8 - 26

#### Amylopectin granules

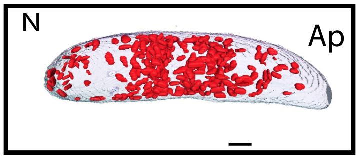


**Volume:**
Mean = 2.66µm3
SD = 1.22µm3
COV = 46%
Range = 1.01µm3 - 5.15µm3

**Number:**
Mean = 196
SD = 63
COV = 32.3%
Range = 86 - 294

#### Acidocalcisomes

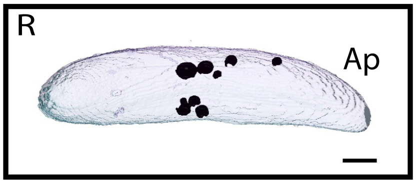


**Volume:**
Mean = 0.32µm3
SD = 0.09µm3
COV = 26.8%
Range = 0.19µm3 - 0.5µm3

**Number:**
Mean = 13
SD = 3.7
COV = 28.9%
Range = 6 - 21
