## Supplementary figures and images for "Cellular electron tomography of the apical complex in the apicomplexan parasite *Eimeria tenella* shows a highly organised gateway for regulated secretion"

### supplemental figure

#### Supplemental Figure 1

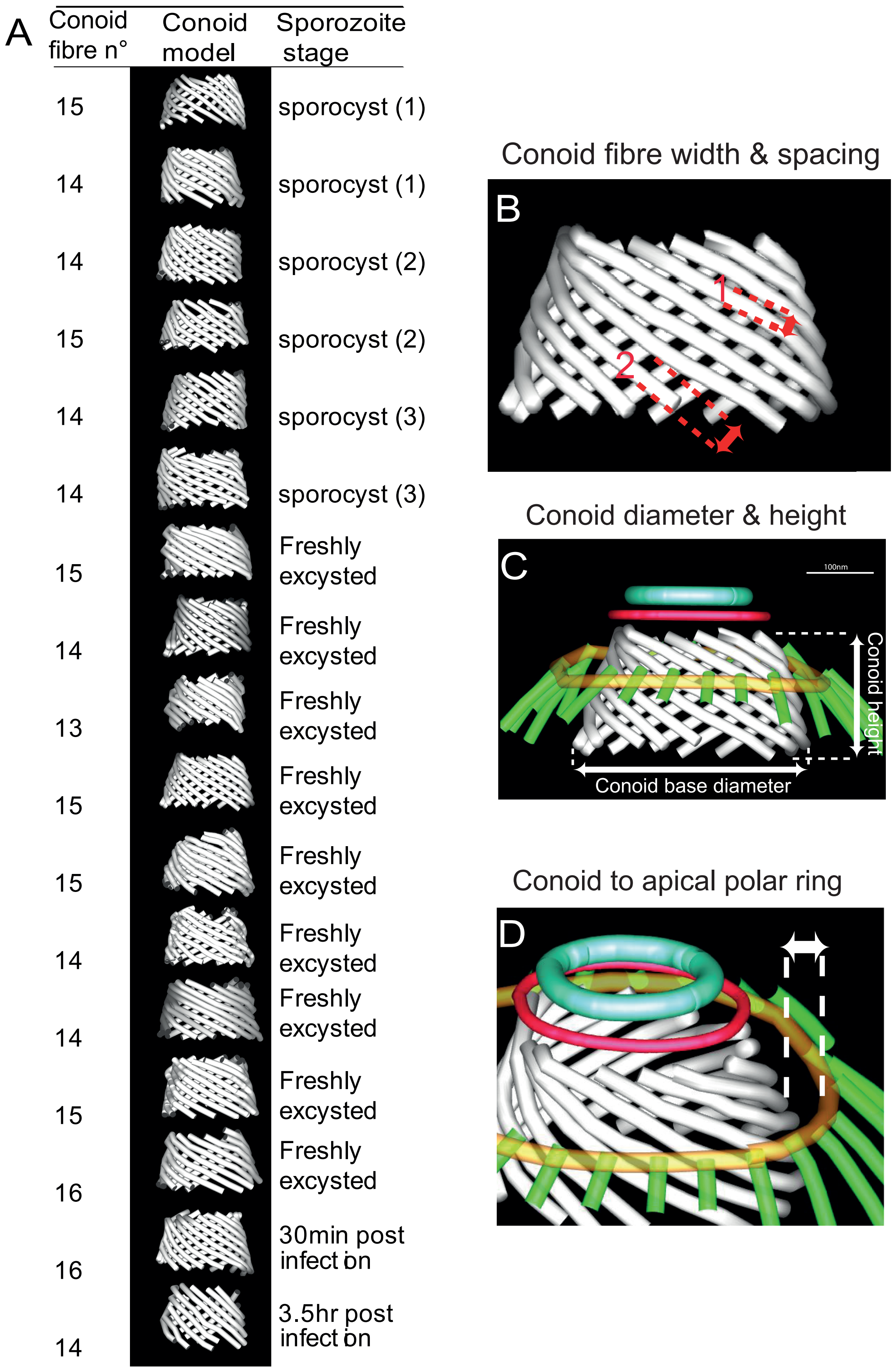

### supplemental figure

Supplemental Figure 2

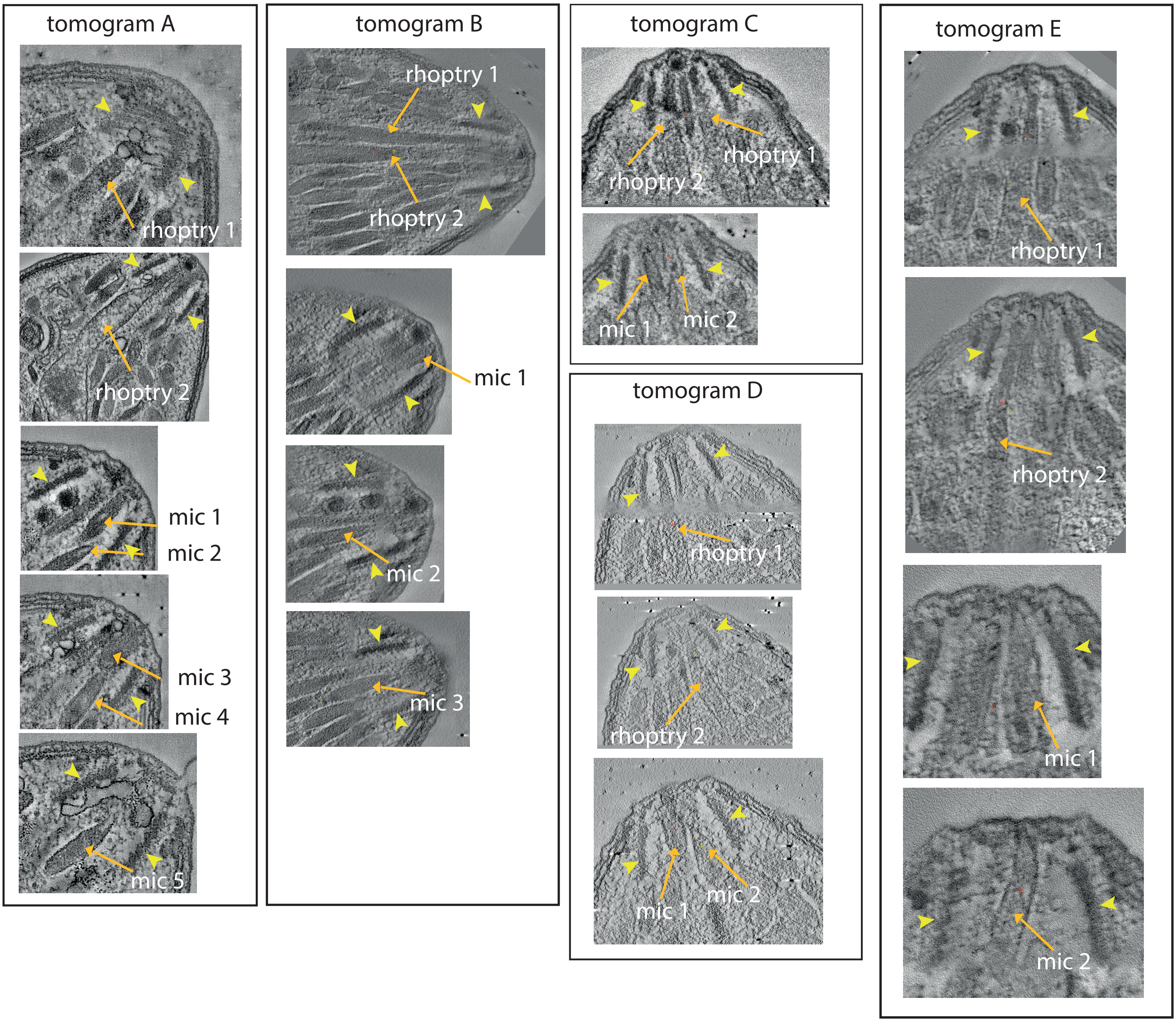
